# Near-infrared carbon nanotube tracking reveals the nanoscale extracellular space around synapses

**DOI:** 10.1101/2021.10.29.466461

**Authors:** Chiara Paviolo, Joana S. Ferreira, Antony Lee, Daniel Hunter, Laurent Groc, Laurent Cognet

**Author notes:** equal contribution (Chiara Paviolo, Joana S. Ferreira). Addresses.

## Abstract

We provide evidence of a local synaptic nano-environment in the brain extracellular space (ECS) lying within 500 nm of postsynaptic densities. To reveal this brain compartment, we developed a correlative imaging approach dedicated to thick brain tissue based on single-particle tracking of individual fluorescent single wall carbon nanotubes (SWCNTs) in living samples and on speckle-based HiLo microscopy of synaptic labels. We show that the extracellular space around synapses bears specific properties in terms of morphology at the nanoscale and inner diffusivity. We finally show that the ECS juxta-synaptic region changes its diffusion parameters in response to neuronal activity, indicating that this nano-environment might play a role in the regulation of brain activity.

---

Neuronal communication in the central nervous system mainly occurs at the level of synapses through the release of neurotransmitters in the synaptic cleft and the activation of postsynaptic receptors. Neurotransmitters can spillover from synapses and act at distance through the so-called “volume transmission”, in which signalling molecules navigate within the brain extra-cellular space (ECS) ^1,2^. Despite technical and molecular advances over the last decades, the local dimensions and architecture of this complex environment are still to be elucidated in identified regions of the living brain. In particular, experimental strategies offering nanoscale resolutions are needed to unveil the narrow and tortuous environment between neural cells^3^.

Changes in the ECS can affect neuronal excitability and signal transmission by altering local ion concentrations in the healthy and diseased brain^4–6^. Compared with free diffusion in an “open space” where molecules move randomly, diffusion in the ECS is critically dependent on the physical and chemical structure of the local microenvironment^7^. Zheng *et al*. investigated the extracellular diffusivity inside the cleft of hippocampal synapses, suggesting reduced diffusion when compared to free medium^8^. Yet, understanding the mechanisms through which the ECS modulates neuronal communication, and particularly in and around synapses, still represents a key challenge in brain research^3^. Because the characteristics of the ECS around synapses remains unknown at the sub-micron scale in native live brain tissue, it is important to establish whether the synaptic ECS environment displays characteristic dimensions and specific diffusional properties.

To tackle this question, we developed a correlative imaging approach based on single-particle tracking of individual fluorescent single wall carbon nanotubes (SWCNTs) in living brain tissues and on speckle-based HiLo microscopy of synaptic labels (Figure 1A). This revealed the existence of a local synaptic nano-environment (which we called “juxta-synaptic” region) lying within 500 nm of postsynaptic densities (PSDs). This local ECS environment is defined by specific properties in terms of morphology at the nanoscale and inner diffusivity. We finally show that the juxta-synaptic region changes its diffusion parameters in response to neuronal activity, indicating that its features might play a role in the regulation of brain activity.

**Figure 1.**
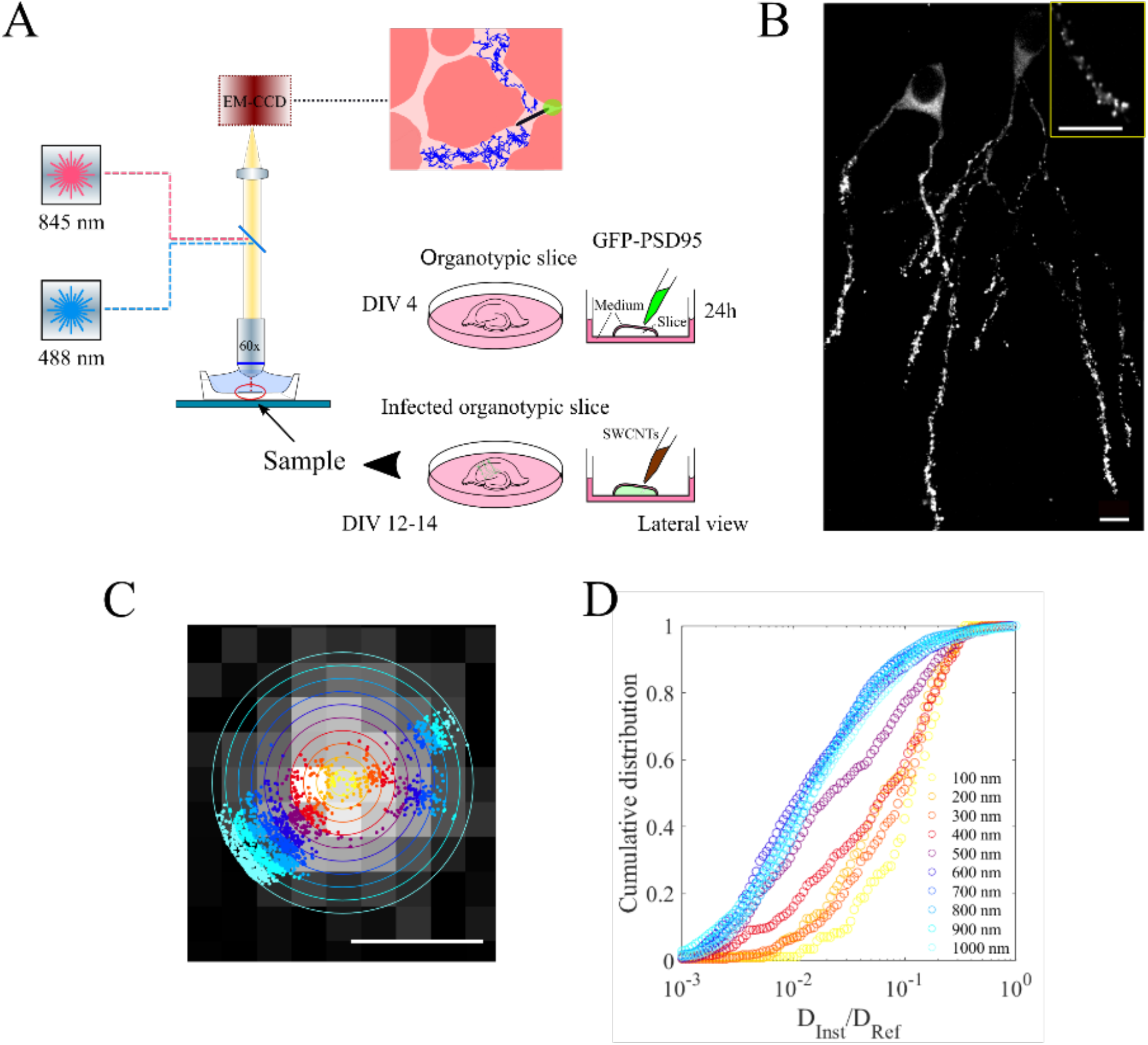
Experimental conditions. A) Schematic of the experimental setup. Rat organotypic cultures were prepared from rat pup brains and cultured for 4 days *in vitro* (DIV) before PSD95 infection. SWCNTs were incubated at DIV 12-14 for 2 hours. NIR imaging of SWCNTs was performed at 845 nm with a 60× immersion objective. GFP-PSD95 expressing neurons were imaged at 488 nm through the same objective. Individual SWCNTs were followed in the ECS of living brains using an EM-CCD camera. B) Image of GFP-PSD95 positive neurons with individual synapses (inset). Scale bars are 10 μm. C) Schematic of the analysis. Individual SWCNT trajectories were analyzed on coronal areas of 100 nm from the synaptic centroids. Scale bar is 1 μm. D) Cumulative distribution functions of diffusivity for untreated samples in each coronal area (31 SWCNTs). A cut-off in the diffusivity values was observed between 400 and 500 nm from the PSD95 centroids.

Near-infrared luminescent SWCNTs are interesting nanoparticles for deep brain tissue microscopy^9^, due to their unique brightness, photostability, and near-infrared emission range in the biological window^10^. SWCNTs also demonstrated capability to locally probe *in situ* chemical species including neurotransmitters^11^. At the single nanotube level, we have recently shown that SWCNTs can access ECS in the brain tissue of young^12^ and adult^13^ rodents, and their diffusion trajectories can reveal ECS local dimensions. Noteworthy, those dimensions are consistent with another ECS imaging approach based on STED microscopy^14^. Using these approaches, ECS remodeling in a neuropathological condition was successfully reported^14,15^. To locally probe the ECS dimension and diffusivity around synapses, the use of SWCNTs in the live brain tissue emerged thus as a promising tool.

In order to identify synaptic areas into cultured hippocampal brain slices, we fluorescently labelled synapses using GFP-PSD95 lentivirus vectors (Figure 1B). PSD95 is one of the most abundant proteins in excitatory synapses and is a common marker of postsynaptic areas. Synaptic imaging was performed in the CA1 region of the hippocampus at depth ranging from 10 to approximately 50 μm (Figure S1A) using a speckle based structured illumination technique known as HiLo microscopy (Sparq module)^16^. HiLo microscopy requires the acquisition of two images to generate good optical sectioning: one structured/speckle image and one uniform illumination image. From HiLo images, GFP positive clusters corresponding to synapses were then identified (Figure 1B inset).

Biocompatible fluorescent SWCNTs were prepared by encapsulation with phospholipid-polyethylene glycol (PL-PEG) molecules. This coating minimizes non-specific adsorption onto biological structures^17^ while preserving SWCNT luminescence brightness for single molecule experiments. Cultured hippocampal slices expressing GFP-PSD95 were incubated with PL-PEG coated SWCNTs and slices were placed onto a NIR single molecule microscope (see material and methods). We focused on (6,5) SWCNTs emitting at 985 nm which are efficiently excited at 845 nm while minimizing light absorption by the tissue^18^. Bright (6,5) SWCNTs were detected at high signal to noise ratio with low autofluorescence coming from biological structures, which constitutes a decisive asset to perform single molecule imaging at required depth. Indeed, investigating ECS structures requires thick brain tissue preparations (generally few hundred micrometers). Luminescent SWCNTs were imaged at video rates to grasp their rapid diffusion within the ECS. Importantly, SWCNT high aspect-ratio and intrinsic rigidity play a decisive role here, slowing down nanotube diffusion in the ECS maze while ensuring high accessibility to nanoscale environments^19^. These are unique features of these bright non photobleaching 1D nanoparticles.

Super-localization analysis of nanotube positions along their trajectories was performed as follows. For each recorded movie frame, we applied a 2D asymmetric Gaussian fitting analysis of the fluorescence profiles to decipher the nanotube centroids with sub-wavelength precision (∼ 50 nm) and nanotube length (long axis of the Gaussian fit). Taking into account the exciton diffusion range that decreases the apparent nanotube length in fluorescence images due to end quenching^20^, the measured nanotube length distribution was centered around ∼600 nm (Figure S1B). This narrow distribution confirms the consistency and reproducibility of the nanotube preparation over 14 independent experiments (3 slices per experiment; 76 analyzed SWCNTs).

In order to explore the ECS environment around synapses, the region around each GFP-PSD95 centroids was segmented by a series of concentric coronal areas of 100 nm widths (Figure 1C). A maximum distance of 1 μm from the synaptic centroids was considered, based on the average synapse density in hippocampal neurons (i.e. around 1 spine per μm of dendrite^21^). An image correlation analysis with SWCNT localizations allowed us to create a distance-to-synapse investigation of the ECS features. We first analysed SWCNT local diffusivity in the different coronal areas. Local diffusivity was defined as the normalized local instantaneous diffusion along trajectories. For this, the two-dimensional mean squared displacement (MSD) was calculated as a function of time intervals along a sliding window of 390 ms for each trajectory. Approximating the MSD as linear at short time intervals, a linear fit of the first 90 ms yielded *D*_*inst*_, the instantaneous diffusion coefficient. The normalized diffusivity was then obtained by calculating *D*_*inst*_/*D*_*ref*_ where *D*_*ref*_ is the calculated diffusion coefficient of carbon nanotube freely diffusing in a fluid having the viscosity *η*_*ref*_ of the cerebrospinal fluid (CSF)^7^ (see Methods).

Figure 1D shows cumulative distributions of local diffusivities measured in different coronal areas around synapses. Clearly, a distance-to-synapse dependent behavior lies within submicron scales. More specifically, a cut-off in the diffusivity is observed between 400 and 500 nm revealing a specific diffusivity behavior around synapses where the particles undergo a 10-fold enhanced diffusivity as compared to away from synapses. In order to unambiguously assess the presence of this specific juxta-synaptic diffusion environment, a series of controls was performed. We simulated SWCNT trajectories assuming Brownian motion and randomly generated synaptic localizations to exclude that the generation of 100 nm width coronal regions might bias apparent diffusivities into reduced coronal areas (Figure S2A-B). Furthermore, we also generated “false” synapses localizations at distance from the measured ones when analyzing the experimental SWCNT trajectories: no specific (enhanced) diffusivities were subsequently generated around “false” synaptic areas (Figure S2C). We thus conclude that a “juxta-synaptic” environment exists up to 500 nm distance from synaptic centroids where diffusivities are larger than in “non-juxta-synaptic” areas (areas between 500 and 1000 nm from GFP-PSD95 centroids).

Based on this partition of the ECS environment definition (juxta- or non-juxta-synaptic), we next ran an extensive characterization of the ECS features pooled into these regions (Figure 2). In addition to information on local diffusivity, the analysis of SWCNT localizations also provides information on the local ECS dimensions applying an analytical approach previously described^13^. In short, SWCNT localizations were fitted to an ellipse over short periods of time (180 ms), where the shorter dimension represents the local ECS dimensions (ξ). Spatial maps of local diffusivities (*D*_*inst*_*/D*_*ref*_) and ECS local dimensions were thus generated (Figure 2A), revealing that SWCNT diffusion is heterogeneous in all ECS areas. Due to the high neuronal density of brain tissues, we cannot exclude that the non-juxta-synaptic region may include synapses from other dendrites of non-infected (non-labelled) neurons, so that non-juxta-synaptic behavior might be contaminated by juxta-synaptic features. In addition, because this work does not super-localize SWCNT in 3D (nor synaptic centroids), the depth of focus of our microscope does not discriminate juxta-synaptic areas along the *z* (optical) axis of the microscope, such that juxta-synaptic regions might also be contaminated by non-juxta-synaptic features. In any case, we found that the juxta-synaptic diffusivity is 6-fold faster than the non-juxta-synaptic region (median__juxta_ = 0.089; median__non-juxta_ = 0.014; *p* < 0.001; Figure 2B) which might in fact represent a lower fold due to the two possible “contaminations” just mentioned.

**Figure 2.**
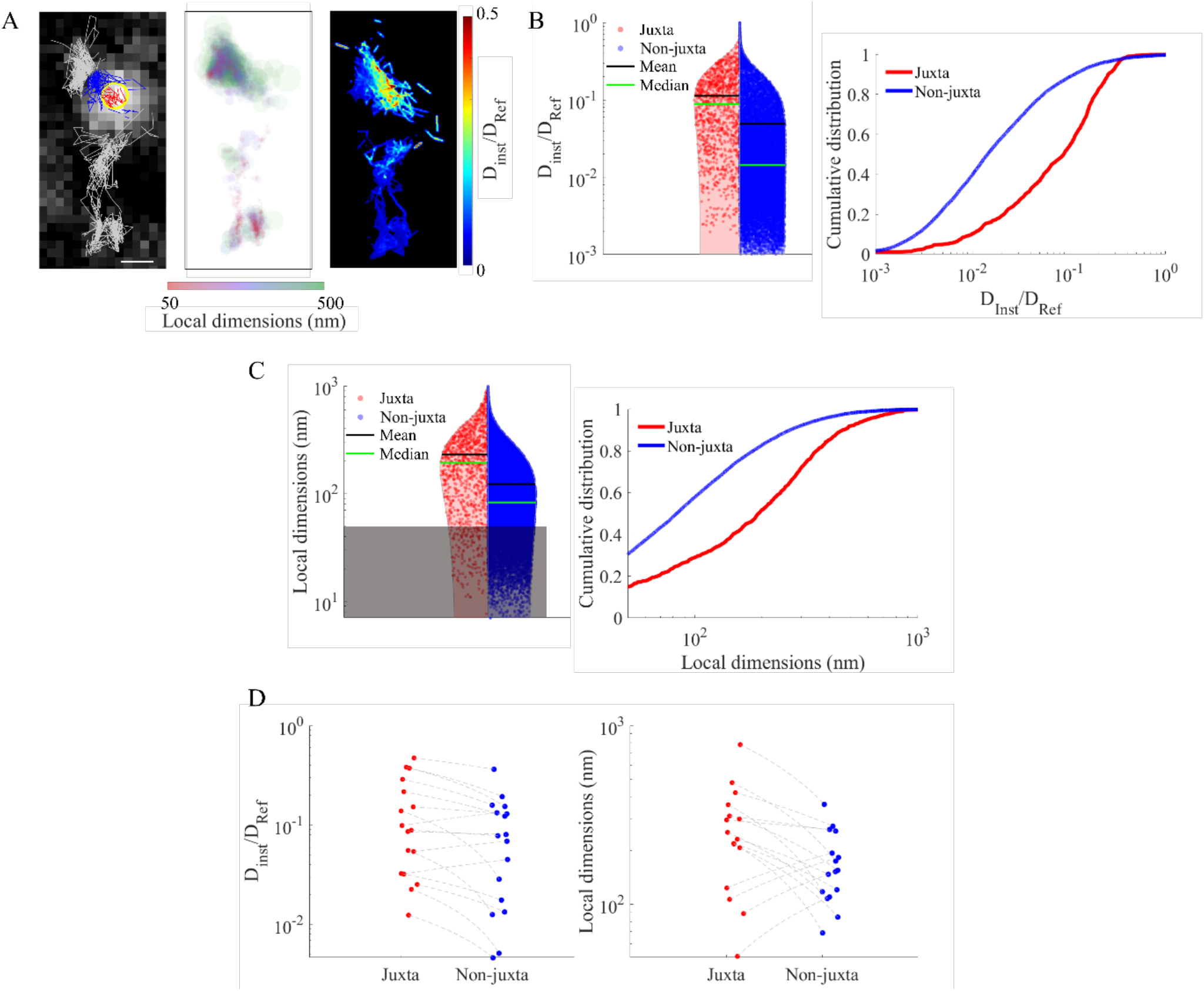
Analysis of untreated organotypic brain slices. A) Examples of individual trajectories. Localizations were divided based on the relative position to a GFP-PSD95 centroid (left panel: juxta-synaptic in red, non-juxta-synaptic in blue). Localizations further than 1 μm are depicted in grey. Super-localization analysis gave further information on local ECS dimensions (central panel) and diffusivity (right panel). Scale bars are 2 μm. B) Violin plot and cumulative distributions of diffusivity and C) local dimensions. Graphs show that the juxta-synaptic nano-environment bears specific local properties (*p* < 0.001). D) Pair analysis of diffusivity and local dimensions in the juxta-synaptic nano-environment (*p* < 0.01).

Figure 2C displays ECS dimension values in juxta- and non-juxta-synaptic domains. Similar to diffusivity, local ECS dimensions were highly heterogeneous, their widths ranging from around 50 nm (limited by the precision of our approach) to well above 1 μm. The vast majority (>70%) of local dimensions in the juxta-synaptic nano-environment were larger than 100 nm. Strikingly the ECS local dimension are significantly larger (∼2-fold) in the juxta-synaptic region as compared to the non-juxta-synaptic ones (median__juxta_ = 193 nm – median__non-juxta_ = 83 nm; *p* < 0.001; Figure 2C). Finally, we performed a paired analysis around each synaptic environment (between juxta- and non-juxta-synaptic regions) for diffusivity and local dimensions (Figure 2D). At the single synapse level, this analysis confirmed that higher diffusivity and larger local ECS dimensions are found in juxta-synaptic environments with respect to the local non-juxta-synaptic nano-environment (*p* < 0.01, Figure 2D).

In general, the broad shape of the cumulative distributions confirmed that the ECS is a highly heterogeneous milieu, where diverse local properties can impose a wide range of diffusivity and dimensional values. Indeed, in contrast to a free medium, diffusion in the brain ECS can be hindered by cell processes, astroglia, macromolecules of the matrix, wall drag, and the presence of charged molecules. In this environment, the diffusivity is also dependent on the hydrodynamic dimension of the diffusing probe, resulting in lower diffusivities for larger objects. Interestingly, median values of diffusivity and local dimensions evaluated at the level of individual juxta-synaptic regions are uncorrelated (Pearson’s *r* = 0.138; Figure S2A), suggesting that *i*) the diffusivity of SWCNTs is mainly influenced by the molecular composition of the space, and *ii*) spatial constrictions of cellular walls are not necessarily the central determinant of ECS diffusion inhomogeneities at the nanoscale near synapses.

As stated above, changes in neuronal activity is likely to alter ECS characteristics^12,14,15^. We thus now question whether these changes also alter ECS diffusivity and morphology in the juxta-synaptic nano-environment. We used two classical protocols to either favour (bicuculline, BIC 40 μM) or decrease (tetrodotoxin, TTX 2 μM) neuronal activity (Figure 3A). As expected, incubation with BIC significantly increased the basal activity whereas TTX suppressed it (Fig. 3B). Additionally, no significant differences were detected on the dimension of the GFP-PSD95 clusters (*p* = 0.1231, Fig. S4).

**Figure 3.**
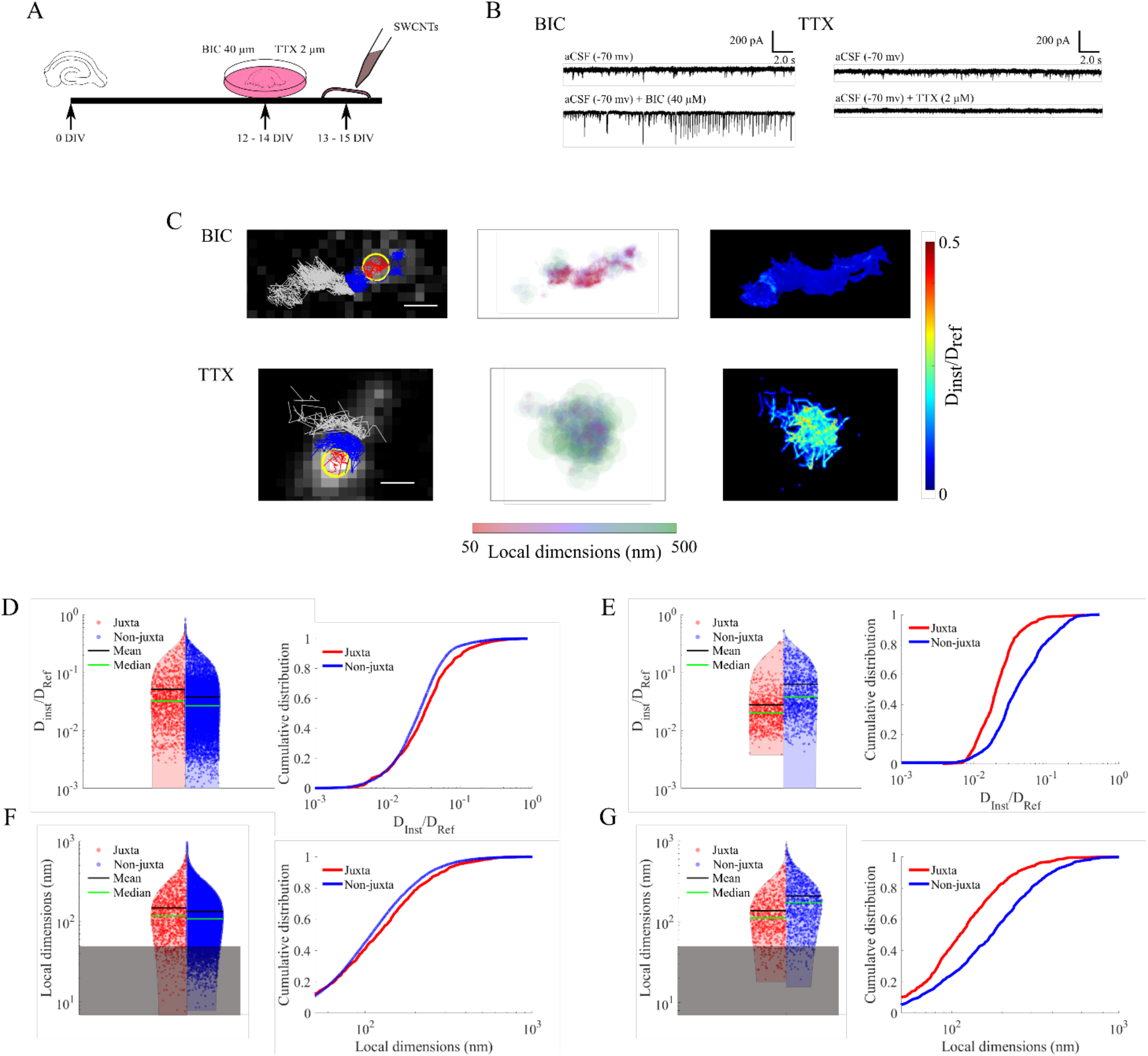
Stimulation of organotypic brain slices. A) Schematic of the stimulation. BIC or TTX treatments were applied for 24 hours prior to SWCNT incubation to respectively block the inhibitory action of GABA_A_ receptors or the sodium channels. B) Representative electrophysiological traces of excitatory postsynaptic currents (EPSCs) from rat organotypic hippocampal slices DIV 12. Top traces recorded in the presence of aCSF only, bottom traces after addition of BIC or TTX in the medium to increase or decrease neuronal activity, respectively. C) Examples of individual trajectories for BIC- and TTX-treated organotypic slices. As for control samples, localizations were divided based on the relative position to a GFP-PSD95 centroid (left panel: juxta-synaptic in red, extra-synaptic in blue). Localizations further than 1 μm are depicted in grey. Super-localization analysis gave further information on local ECS dimensions (central panel) and diffusivity (right panel). Scale bars are 2 μm. D) Violin plot and cumulative distributions of diffusivity for BIC-treated samples. Graphs show that BIC uniformed the synaptic microenvironment. E) Violin plot and cumulative distributions of diffusivity for TTX-treated samples, showing a significant slowing-down of diffusivity in the juxta-synaptic nano-environment. F) Violin plot and cumulative distributions of local dimensions for BIC-treated samples, confirming the uniformity of the environment. G) Violin plot and cumulative distributions of local dimensions for TTX-treated samples. For all graphs, the juxta-synaptic localizations are marked in red, while the non-juxta-synaptic ones in blue. The grey boxes represent the localization precision of our analysis.

Figure 3C shows examples of trajectories of SWCNTs in hippocampal tissues exposed to BIC or TTX. Similar to untreated samples, we partitioned nanotube localizations based on their juxta- or non-juxta-synaptic position. We observed that modulation of neuronal activity is accompanied by changes of the ECS local environment around synapses. More precisely, comparing ECS domains in each condition, we found that for BIC-treated samples the difference in diffusivity between regions near or distant from the GFP-PSD95 centroid became less pronounced (median__juxta_BIC_ = 0.032 – median__non-juxta_ = 0.027; Figure 2D), whereas TTX yielded significant lower values of diffusivity in the juxta-synaptic environment compared to non-juxta-synaptic spaces (median__juxta_ = 0.020 – median__non-juxta_ = 0.038; Figure 2E). A similar alteration/modification was detected after the analysis of local dimensions in the juxta- and non-juxta-synaptic regions (median__juxta_BIC_ = 118 nm – median__non-juxta_BIC_ = 108 nm; Figure 2F - median__juxta_TTX_ = 112 nm – median__non-juxta_TTX_ = 174 nm; Figure 2G).

We next focus on the juxta synaptic region to compare the different conditions (Fig. 4). SWCNT diffusivity was slowed down and ECS local dimensions were shrank in both BIC and TTX treatments when compared to control conditions (Figure 4A-B, *p* < 0.001). This observation suggests that the neuronal network can accommodate to neuronal activity changes (either increase or blockade) through ECS regulation. Finally, Figure 4 also indicates that in the juxta-synaptic region distributions of local diffusivity and dimensions of treated samples are more monodisperse than control conditions, suggesting that BIC and TTX treatments standardized the juxta-synaptic environment.

**Figure 4.**
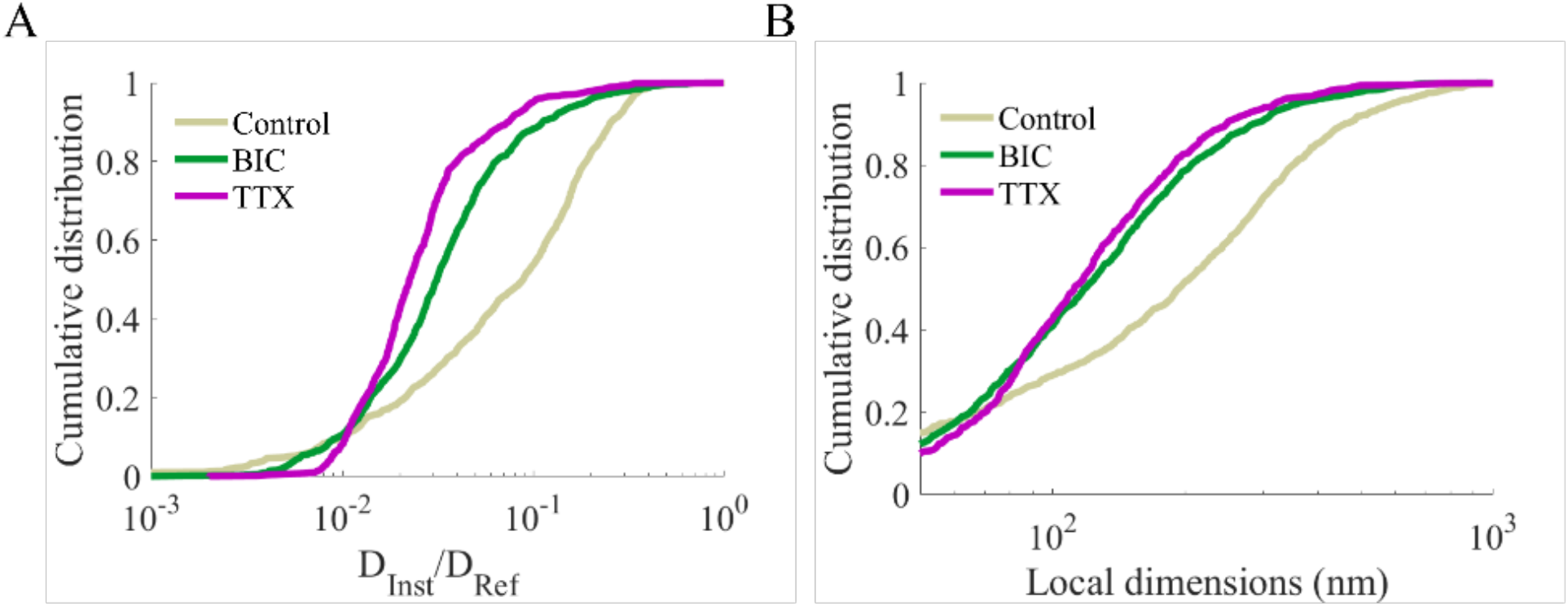
Comparison of cumulative distributions of A) local diffusivity and B) local dimensions for control (grey) BIC- (green), and TTX-treated (purple) organotypic slices. When compared to control samples, both BIC and TTX slowed down the SWCNT diffusivity in the juxta-synaptic nano-environment and narrowed down the local dimensions (*p* < 0.001). Local diffusivity in TTX-treated samples was significantly slower than in BIC-incubated tissues (*p* < 0.001), but the local dimensions remained comparable between the two conditions.

A significant and global decrease in the ECS volume fraction and an increase in diffusion barriers have been reported during neuronal activity and pathological states^22^. These changes were related to cell swelling, cell loss, astrogliosis, rearrangement of neuronal and astrocytic processes, and changes in the extracellular matrix. Plastic changes in ECS volume, tortuosity, and anisotropy can also affect the neuronal-glial communication, the spatial relation of glial processes toward synapses, glutamate or GABA ‘spillover’, and/or synaptic crosstalk^5,6^. Here, TTX-treated samples have slower diffusivity than the ones incubated with BIC (*p* < 0.001), but the local dimensions remain comparable between conditions, suggesting that the differences between BIC and TTX relies on the chemical modifications of the juxta-synaptic region. This is further supported by the correlation analysis evaluated for individual GFP-PSD95 positive clusters, which revealed a higher correlation between local diffusivity and dimensions in the juxta-synaptic region for BIC-with respect to TTX-treated samples (Figure S3).

Altogether, our study unveiled the existence of a juxta-synaptic ECS nano-environment within 500 nm from a synaptic centre. This observation was possible by correlating the dynamics and super-localization of NIR-emitting carbon nanotube with HiLo microscopy of labelled synapses in live brain slices. Increasing or decreasing synaptic activity specifically modified the ECS diffusion and morphological parameters in the juxta-synaptic region. Such a regulation of the ECS nano-environment around synapses would strongly influence the diffusion of neurotransmitters and modulators in the brain tissue, impacting neuronal network physiology and pathology.

## ACKNOWLEDGEMENTS

This work was performed with financial support from by the European Research Council Synergy grant (951294), Agence Nationale de la Recherche (ANR-15-CE16-0004-03), and the France-BioImaging National Infrastructure (ANR-10-INBS-04-01). C.P. acknowledges funding from EU’s Horizon 2020 research and innovation program under the Marie Skłodowska-Curie grant No 793296. A.L. acknowledges support from the Fondation ARC pour la recherche sur le cancer. We thank the Bordeaux Imaging Center, a service unit of the CNRS-INSERM and Bordeaux University, the IINS Cell Biology Facility for organotypic culture slices preparation, in particular the help of Emeline Verdier, Delphine Bouchet Tessier and Constance Manso. We thank Christophe Mulle and Severine Deforges for providing the GFP-PSD95 lentivirus.

## REFERENCES

(1) Nicholson, C.; Syková, E. Extracellular Space Structure Revealed by Diffusion Analysis. Trends Neurosci. 1998, 21 (5), 207–215.

(2) Rusakov, D. A.; Min, M.-Y.; Skibo, G. G.; Savchenko, L. P.; Stewart, M. G.; Kullmann, D. M. Role of the Synaptic Microenvironment in Functional Modification of Synaptic Transmission. Neurophysiology 1999, 31 (2), 79–81.

(3) Nicholson, C.; Hrabětová, S. Brain Extracellular Space: The Final Frontier of Neuroscience. Biophys. J. 2017, 113 (10), 2133–2142.

(4) Dietzel, I.; Heinemann, U.; Hofmeier, G.; Lux, H. D. Transient Changes in the Size of the Extracellular Space in the Sensorimotor Cortex of Cats in Relation to Stimulus-Induced Changes in Potassium Concentration. Exp. Brain. Res. 1980, 40 (4), 432–439.

(5) Slais, K.; Vorisek, I.; Zoremba, N.; Homola, A.; Dmytrenko, L.; Sykova, E. Brain Metabolism and Diffusion in the Rat Cerebral Cortex during Pilocarpine-Induced Status Epilepticus. Exp. Neurol. 2008, 209 (1), 145–154.

(6) Colbourn, R.; Naik, A.; Hrabětová, S. ECS Dynamism and Its Influence on Neuronal Excitability and Seizures. Neurochem. Res. 2019, 44 (5), 1020–1036.

(7) Syková, E.; Nicholson, C. Diffusion in Brain Extracellular Space. Physiol. Rev. 2008, 88 (4), 1277–1340.

(8) Zheng, K.; Jensen, T. P.; Savtchenko, L. P.; Levitt, J. A.; Suhling, K.; Rusakov, D. A. Nanoscale Diffusion in the Synaptic Cleft and beyond Measured with Time-Resolved Fluorescence Anisotropy Imaging. Sci. Rep. 2017, 7, 42022.

(9) Paviolo, C.; Cognet, L. Near-Infrared Nanoscopy with Carbon-Based Nanoparticles for the Exploration of the Brain Extracellular Space. Neurobiology of Disease 2021, 153, 105328. https://doi.org/10.1016/j.nbd.2021.105328.

(10) Welsher, K.; Liu, Z.; Sherlock, S. P.; Robinson, J. T.; Chen, Z.; Daranciang, D.; Dai, H. A Route to Brightly Fluorescent Carbon Nanotubes for Near-Infrared Imaging in Mice. Nat. Nanotechnol. 2009, 4 (11), 773–780.

(11) Kruss, S.; Landry, M. P.; Vander Ende, E.; Lima, B. M. A.; Reuel, N. F.; Zhang, J.; Nelson, J.; Mu, B.; Hilmer, A.; Strano, M. Neurotransmitter Detection Using Corona Phase Molecular Recognition on Fluorescent Single-Walled Carbon Nanotube Sensors. J. Am. Chem. Soc. 2014, 136 (2), 713–724.

(12) Godin, A. G.; Varela, J. A.; Gao, Z.; Danné, N.; Dupuis, J. P.; Lounis, B.; Groc, L.; Cognet, L. Single-Nanotube Tracking Reveals the Nanoscale Organization of the Extracellular Space in the Live Brain. Nat. Nanotechnol. 2017, 12 (3), 238–243.

(13) Paviolo, C.; Soria, F. N.; Ferreira, J. S.; Lee, A.; Groc, L.; Bezard, E.; Cognet, L. Nanoscale Exploration of the Extracellular Space in the Live Brain by Combining Single Carbon Nanotube Tracking and Super-Resolution Imaging Analysis. Methods 2020, 174, 91–99.

(14) Tønnesen, J.; Inavalli, V. V. G. K.; Nägerl, U. V. Super-Resolution Imaging of the Extracellular Space in Living Brain Tissue. Cell 2018, 172 (5), 1108-1121.e15.

(15) Soria, F. N.; Paviolo, C.; Doudnikoff, E.; Arotcarena, M.-L.; Lee, A.; Danné, N.; Mandal, A. K.; Gosset, P.; Dehay, B.; Groc, L.; Cognet, L.; Bezard, E. Synucleinopathy Alters Nanoscale Organization and Diffusion in the Brain Extracellular Space through Hyaluronan Remodeling. Nat. Commun. 2020, 11 (1), 3440.

(16) Lim, D.; Ford, T. N.; Chu, K. K.; Mertz, J. Optically Sectioned in Vivo Imaging with Speckle Illumination HiLo Microscopy. J. Biomed. Opt. 2011, 16 (1), 016014-016014–016018.

(17) Gao, Z.; Danné, N.; Godin, A. G.; Lounis, B.; Cognet, L. Evaluation of Different Single-Walled Carbon Nanotube Surface Coatings for Single-Particle Tracking Applications in Biological Environments. Nanomaterials 2017, 7 (11), 393.

(18) Danné, N.; Godin, A. G.; Gao, Z.; Varela, J. A.; Groc, L.; Lounis, B.; Cognet, L. Comparative Analysis of Photoluminescence and Upconversion Emission from Individual Carbon Nanotubes for Bioimaging Applications. ACS Photonics 2018, 5 (2), 359–364.

(19) Fakhri, N.; MacKintosh, F. C.; Lounis, B.; Cognet, L.; Pasquali, M. Brownian Motion of Stiff Filaments in a Crowded Environment. Science 2010, 330 (6012), 1804–1807.

(20) Oudjedi, L.; Parra-Vasquez, A. N. G.; Godin, A. G.; Cognet, L.; Lounis, B. Metrological Investigation of the (6,5) Carbon Nanotube Absorption Cross Section. J. Phys. Chem. Lett. 2013, 4 (9), 1460–1464.

(21) De Simoni, A.; Griesinger, C. B.; Edwards, F. A. Development of Rat CA1 Neurones in Acute versus Organotypic Slices: Role of Experience in Synaptic Morphology and Activity. J. Physiol. 2003, 550 (Pt 1), 135–147.

(22) Vargová, L.; Syková, E. Astrocytes and Extracellular Matrix in Extrasynaptic Volume Transmission. Philos. Trans. R. Soc. Lond., B, Biol. Sci. 2014, 369 (1654), 20130608.

