## Supplemental material for "Near-infrared carbon nanotube tracking reveals the nanoscale extracellular space around synapses"

[#] equal contribution

### MATERIALS AND METHODS

*Rat organotypic slice preparation*

Organotypic slice cultures were prepared as previously described in Paviolo *et al.*[1]. Hippocampal slices (350 μm) were obtained from postnatal day 5 to 7 Sprague-Dawley (Janvier Labs) rats using a McIlwain tissue chopper and then placed in dissection medium containing (in mM): 175 sucrose, 25 D-glucose, 50 NaCl, 0.5 $CaCl_2$, 2.5 KCl, 0.66 $KH_2PO_4$, 2 $MgCl_2$, 0.28 $MgSO_4$-7$H_2O$, 0.85 $Na_2HPO_4$-12$H_2O$, 2.7 $NaHCO_3$, 0.4 HEPES, 2×10–5% phenol red, pH 7.3. Slices were kept at 4°C until they were transferred to hydrophilic polytetrafluoroethylene (FHLC) membranes (Millipore) set on Millicell Cell Culture Inserts (Millipore), containing pre-warmed culture medium (50% Basal Medium Eagle, 25% Hank's balanced salt solution, 25% horse serum, 0.45% D-glucose, and 1mM L-glutamine) and cultured for up to 14 days at 35°C / 5% $CO_2$ and the medium replaced every 2 to 3 days.

*Lentivirus construction and slices infection*

Lentivirus transfer vector construct (FHUG+W) expressing PSD95 tagged with GFP[2], was produced at the Vectorology Platform (INSERM US 005 – CNRS 3427 – TBMCore, Université de Bordeaux, France) by transfection with a 3 viral vector system — mock with the PSD95-GFP insert, pCMV-Δ8-9 (encapsulation plasmid), and VSV-G (cDNA encoding the envelope glycoprotein of vesicular stomatitis virus) — in FT-HEK293 cells. Lentivirus supernatants were concentrated by centrifugation concentration filter (Centricon) and a final titter of $2.14^8$ virus particles/ml was obtained. 1 μl of the concentrated lentivirus was added to FHLC membranes, cut to the slice size, and immediately inverted over the slice at 4 days of culture *in vitro* (DIV 4) for 24h. After FHLC membrane removal, medium was replaced with fresh one.

*Confocal images of infected slices*

Images of GFP-PSD95 expressing neurons in organotypic slices at DIV 12 were taken in an inverted Leica DMI 6000 microscope (Leica Microsystems, Wetzlar, Germany) equipped with a confocal head Yokogawa CSU-X1 (Yokogawa Electric Corporation, Tokyo, Japan), a sensitive Quantem camera

(Photometrics, Tucson, USA), using a HCX PL APO CS 63× oil 1.32 NA objective and a 491 nm diode laser. This system was controlled by MetaMorph software (Molecular Devices, Sunnyvale, USA).

*SWCNT preparation*

SWCNTs were prepared as previously described with minor modifications[1]. Briefly, 1 mg of HiPco synthesized carbon nanotubes (from Rice University) was suspended with 50 mg of monofunctional phospholipid-polyethylene glycol (PL-PEG) (#mPEG-DSPE-5000, Laysan Bio) in 10 ml of deuterium oxide (Sigma Aldrich). To individually disperse the nanotubes, a 15 min homogenization at 19,000 rpm followed by an 8 min tip sonication at 20W were applied to the solution. SWCNT bundles and impurities were further precipitated by centrifugation at 3,000 rpm for 60 min. 70–80% of the supernatant was then collected and stored at 4 °C.

*Slice stimulation and SWCNT incubation*

For GFP-PSD95-infected slice stimulation, 40 µM of (-)-Bicuculline methochloride (BIC, TOCRIS) or 2 µM of tetrodotoxin (TTX, TOCRIS) were applied for 24 hours to respectively block the inhibitory action of $GABA_A$ receptors or the sodium channels at DIV 12-14. After the incubation time, slices returned in fresh culture medium. SWCNTs were incubated in the cultures 2 hours prior imaging. 3 µl of SWCNT solution was mixed with 100 µl of culture medium and incubated with the GFP-PSD95-infected slices at 35 °C / 5% $CO_2$. Slices were imaged for up to 1 h in HEPES-based artificial cerebrospinal fluid (HEPES-aCSF) containing (in mM): 130 NaCl, 2.5 KCl, 2.2 $CaCl_2$, 1.5 $MgCl_2$, 10 HEPES, and 10 D-glucose.

*Electrophysiology of BIC and TTX treated slices*

For electrophysiological experiments, whole-cell voltage-clamp recordings were taken from CA1 pyramidal cells in organotypic hippocampal slice cultures at DIV 12. Slices were transferred to the recording chamber, perfused with 32°C artificial cerebrospinal fluid (aCSF) composed of (in mM): 126 NaCl, 3.5 KCl, 2 CaCl2, 1.3 MgCl2, 1.2 NaH2PO4, 25 NaHCO3 and 12.1 glucose. For pharmacological manipulations, aCSF perfusion was supplemented with bicuculline (BIC, 40µM) to block GABARmediated inhibition; or with tetrodotoxin (TTX, 2μM) to block action potential-driven synaptic communication. Recording pipettes with a 5-6 MΩ resistance were filled with intracellular solution (in mM): 134 caesium methanesulfonate, 10 HEPES, 0.5 EGTA, 4 Mg-ATP, 0.3 Na-GTP and 4 NaCl. After achieving whole-cell configuration, cells were voltage-clamped at -70mV to record excitatory postsynaptic currents, using a MultiClamp™ 700B amplifier (Molecular Devices), and digitised by Axon Digidata 1550B (Molecular Devices). Baseline recordings were first obtained in the absence of BIC or TTX. Current traces were prepared using Clampfit (v10.7) software.

*Correlative imaging of SWCNT and GFP-PSD95*

Imaging was performed on a customized epifluorescent microscope (Nikon) equipped with a water-cooled EM-CCD camera (ProEM-HS, Princeton Instrument) and a Sparq system (Bliq Photonics). A standard 4× objective (NA 0.1, Nikon) was initially used to check the CA1 position in the hippocampal slice. A 845 nm laser was used to excite individual (6,5) SWCNTs at their phonon sideband ($\lambda_{exc}$ = 845 nm / $\lambda_{em}$ = 986 nm) with a circular polarized excitation. Images were collected using a water immersion 60× objective (NA 1.0, Nikon) using an exposure time of 30 ms. GFP-PSD95 visualization was performed using a 488 nm laser (Coherent) coupled with the Sparq module. Visible images were collected at the end of each SWCNT recording. Recording depth was measured using white light illumination.

*Analysis of GFP-PSD95 cluster areas*

GFP-PSD95 positive clusters selected for analysis were subjected to an intensity threshold to define the cluster, followed by the measurement of the selected area with ImageJ version 1.53j.

*Super-localization analysis*

Super-localization of the SWCNT centroids was obtained by fitting the NIR images with two-dimensional asymmetric Gaussian functions having arbitrary orientations. Three consecutive images were averaged for each fit to improve the localization precision (~50 nm in water). Eventual drifts were corrected using an immobile SWCNT in the field of view. SWCNT coordinates were then interconnected to reconstruct nanotube trajectories. GFP-PSD95 centroids were also estimated by fitting the visible images with two-dimensional asymmetric Gaussian functions. The distance between the SWCNT and the GFP-PSD95 centroids gave the relative position of the nanotube to a synapse. Diffusivity and local dimension analysis were computed on individual trajectories in coronal areas of 100 nm from the synaptic centroids.

Local ECS dimensions were estimated as previously described. Briefly, SWCNT localizations (typically 5000 points) were fitted to an ellipse for time windows of 180 ms. The shorter dimension of the ellipse was then used to define the local ECS dimensions (ξ) as described in [1].

*Local relative diffusivity*

Analysis of individual diffusing SWCNTs in the ECS of live tissues was performed as follows. For each trajectory, the SWCNT length was estimated using the distribution of the longest axis of the 2D asymmetric Gaussian fits for negligible SWCNT movements (displacements between consecutive images < 40 nm) corrected by the point-spread function of the microscope and the exciton diffusion length[3].

The instantaneous mean square displacement (*MSD*) was then calculated for each trajectory as a function of time intervals $\Delta t$ using sliding windows of 390 ms. For short time delays (90 ms), the two-dimensional MSD can be approximated by a linear slope,

$$MSD(t) = 4D_{inst}\Delta t$$

therefore allowing the definition of $D_{inst}$, the instantaneous diffusion coefficient. The localisation precision was estimated from the intercept of the MSD axis at $t$=0 (~ 50 nm). The local relative diffusivity was defined as the ratio between $D_{inst}$ and the value of free diffusion ($D_{ref}$) that the considered SWCNT would have in a fluid bearing the viscosity of the cerebrospinal fluid ($\eta_{ref}$):

$$D_{ref} = \frac{3k_B T ln(2\varphi)}{8\pi\eta_{ref}L}$$

where $k_B$ is the Boltzmann constant, $T$ is the temperature, $\varphi$ is the SWCNT aspect ratio and $L$ is the nanotube length. For visualization purposes, the spatial diffusivity maps were convoluted with a 2D Gaussian of 50 nm full width at half maxima.

*Statistics*

A total of 76 SWCNTs has been analysed in this study (31 for control experiments, $n = 6$; 31 for BIC-treated samples, $n = 6$; 14 for TTX experiments, $n = 4$). Image analysis was performed using custom MATLAB (MathWorks) and Python scripts unless specified otherwise. Statistical analyses were performed in GraphPad Prism 6.01 software or MATLAB. Empirical cumulative distributions were analysed by Kolmogorov–Smirnov (KS) test. Comparison between juxta- and non-juxta-synaptic local dimensions of individual GFP-PSD95 positive clusters was analysed by Paired *t*-test and diffusivity was analysed by Wilcoxon matched-pairs signed rank test. GFP-PSD95 area comparison was analysed by 1-way ANOVA. Statistics for linear correlation was evaluated with Pearson's *r*, using a Student's *t* distribution for the transformation of the correlation. For all statistical tests, the level of significance was set to $p < 0.05$. Discrete data are represented as mean ± standard error of the mean (SEM).

**TABLES**

**Table 1**

| | **Control** | **BIC** | **TTX** |
|---|---|---|---|
| **# SWCNTs** | 31 (*n* = 6) | 31 (*n* = 4) | 14 (*n* = 4) |
| **# GFP-PSD95s** | 17 | 18 | 9 |

Summary of the experimental repetitions. The number of SWCNTs denote the number of trajectories analysed. The numbers of GFP-PSD95 positive clusters represent the quantity of SWCNTs that entered the juxta-synaptic nano-environment. Qualitatively, in TTX-treated samples, SWCNT were more constrained.

**FIGURES**

**Figure S1**

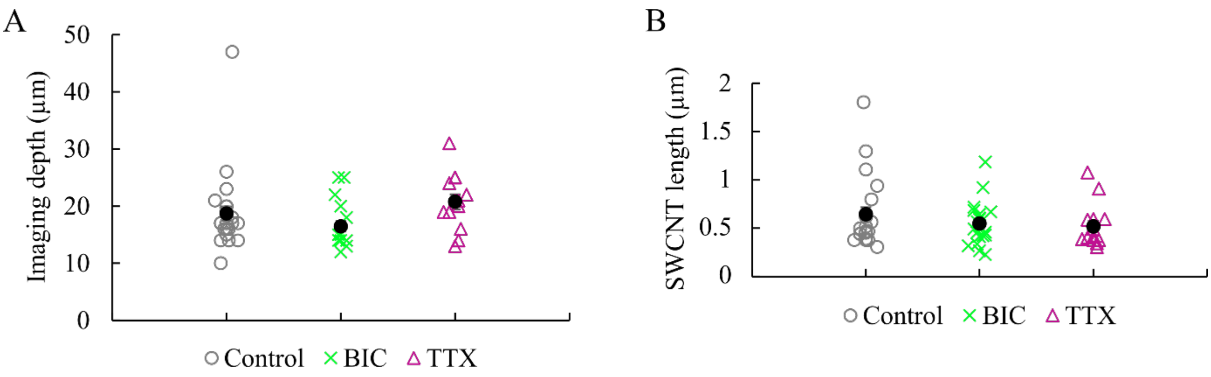


Summary of the experimental conditions in terms of imaging depth (A) and SWCNT length (B) in control, BIC, and TTX conditions.

**Figure S2**

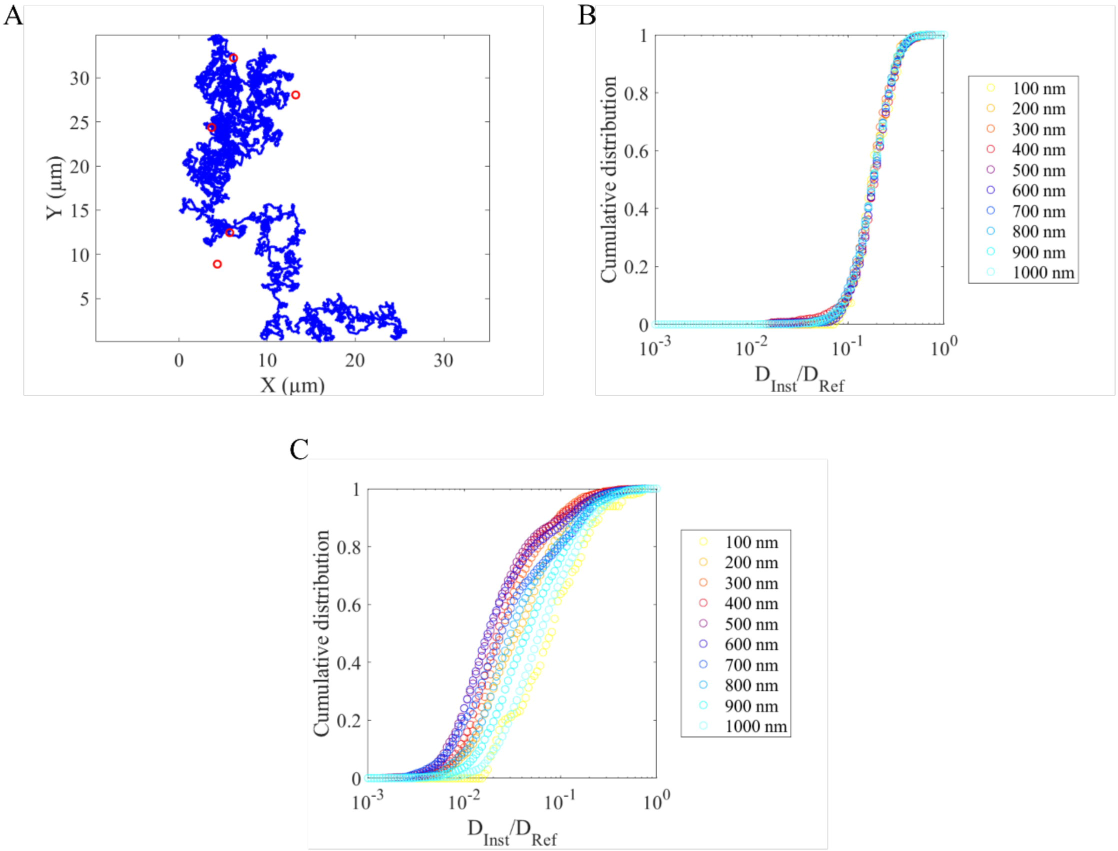


Negative controls for coronal areas. A) Simulated Brownian motion and randomly generated synaptic localization (in red, radius = 400 nm). B) Cumulative distribution functions of diffusivity in each coronal area for the simulated trajectories (N = 30). C) Cumulative distribution functions of diffusivity in each coronal area with randomly generated synapses. Data were generated from experimental SWCNT trajectories. No monotonous distance-to-synapse dependency is found.

**Figure S3**

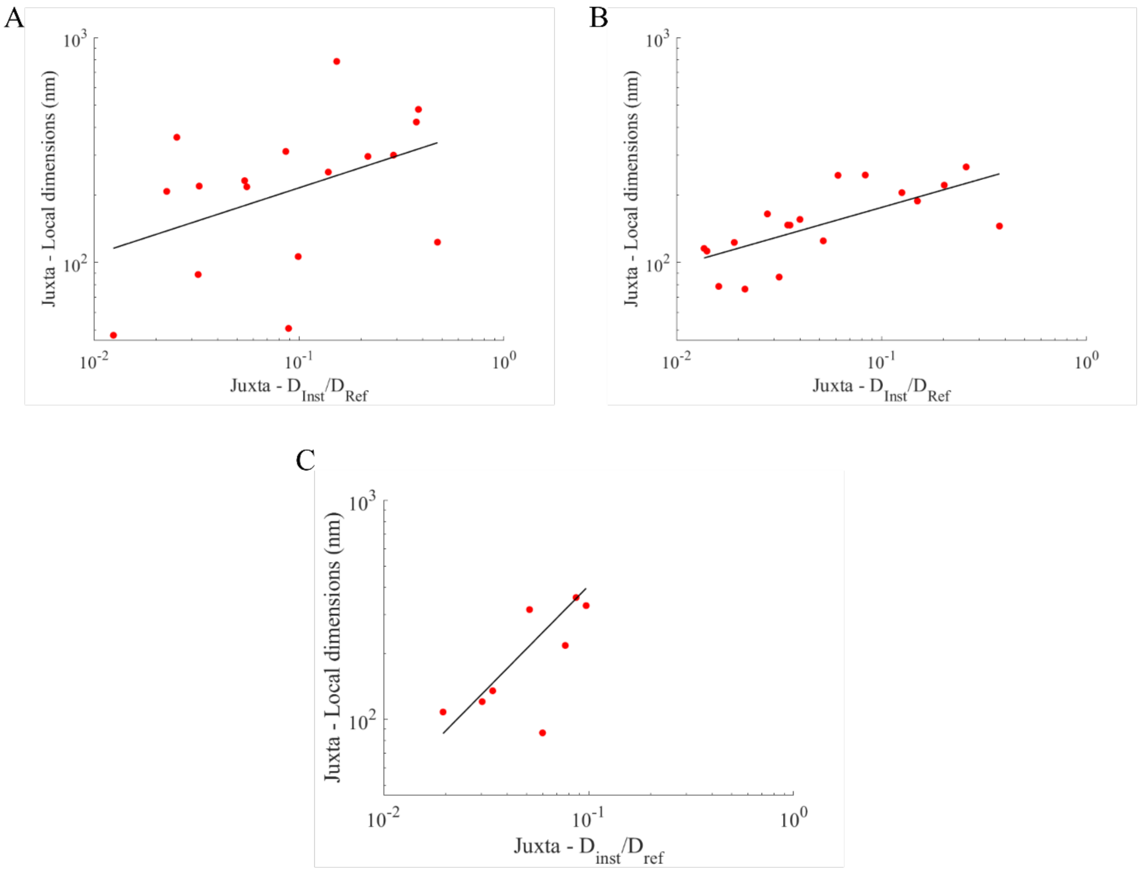


Correlation of local dimensions and diffusivity in the juxta-synaptic nano-environment on individual GFP-PSD95 clusters for control (A), BIC (B), and TTX (C) conditions. Median values of the parameters only showed a low or mild correlation for control and TTX samples (Pearson's $r$ 0.375 and 0.533, respectively), suggesting that the diffusivity of SWCNTs in these environments was mainly influenced by the molecular composition of the space. Analysis of BIC-treated samples revealed a higher correlation (Pearson's $r$ 0.656) between local dimensions and diffusivity.

**Figure S4**

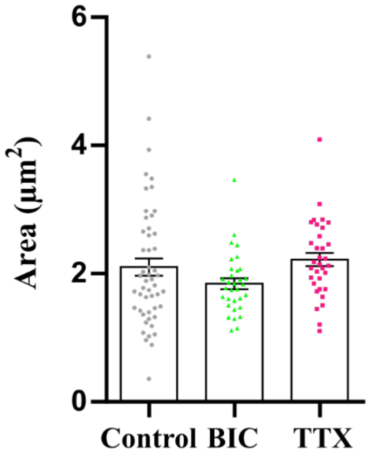


Comparison between GFP-PSD95 cluster areas in control, BIC, and TTX conditions. Treatment did not significantly change the size of GFP-PSD95 positive clusters, used for the corona definition.

### REFERENCES


(1) Paviolo, C.; Soria, F. N.; Ferreira, J. S.; Lee, A.; Groc, L.; Bezard, E.; Cognet, L. Nanoscale Exploration of the Extracellular Space in the Live Brain by Combining Single Carbon Nanotube Tracking and Super-Resolution Imaging Analysis. *Methods* **2020**, *174*, 91–99.

(2) Porras, G.; Berthet, A.; Dehay, B.; Li, Q.; Ladepeche, L.; Normand, E.; Dovero, S.; Martinez, A.; Doudnikoff, E.; Martin-Négrier, M.-L.; Chuan, Q.; Bloch, B.; Choquet, D.; Boué-Grabot, E.; Groc, L.; Bezard, E. PSD-95 Expression Controls L-DOPA Dyskinesia through Dopamine D1 Receptor Trafficking. *J. Clin. Invest.* **2012**, *122* (11), 3977–3989.

(3) Oudjedi, L.; Parra-Vasquez, A. N. G.; Godin, A. G.; Cognet, L.; Lounis, B. Metrological Investigation of the (6,5) Carbon Nanotube Absorption Cross Section. *J. Phys. Chem. Lett.* **2013**, *4* (9), 1460–1464.
